# netneurotools: a trainee-oriented approach to network neuroscience

**DOI:** 10.1101/2025.09.09.675160

**Authors:** Zhen-Qi Liu, Vincent Bazinet, Justine Y. Hansen, Filip Milisav, Andrea I. Luppi, Eric G. Ceballos, Asa Farahani, Laura E. Suarez, Golia Shafiei, Ross D. Markello, Bratislav Misic

## Abstract

Brain imaging is an increasingly inter-disciplinary field, encompassing multiple data types and multiple analytic traditions. Projects typically involve many moving parts, such as building customized preprocessing pipelines, transforming between data formats, preparing datasets for analysis, and ultimately displaying results. The field is conventionally built on highly specialized software packages that solve these individual challenges well, but are not necessarily designed to be interoperable. Trainees new to the field are therefore often left to come up with isolated heuristics and workarounds to complete a project. Here we present a way to navigate the increasingly complex informatics ecosystem of brain imaging. `netneurotools` is our lab’s internal Python toolkit that has been continuously developed and maintained by the lab’s trainees. The philosophy of the toolkit is that it should be the Swiss army knife of the lab: functions and routines that we often use but that are not part of any established pipeline or package. Since its inception, the toolkit has been open and welcomes contribution from neuroscientists across the globe. `netneurotools` presents a necessary counterweight to out-of-the-box software packages and highlights the importance of smaller, ad hoc functions for implementing projects. By opening a window into the inner workings of a lab, `netneurotools` also presents an opportunity to begin a new type of discourse among groups and establish tangible links within the community.

## INTRODUCTION

Modern neuroimaging research requires fluency in many domains, from computing to analytics. Over the past twenty years, numerous software packages have been developed to expedite acquisition, processing, analysis, and visualization of imaging data. Software packages that complement current conceptual, technological, and analytic trends are more commonly used, while others become obsolete. The most enduring packages are often highly specialized, leaving significant daylight between packages that has to be filled by trainees working on a project.

How does a new trainee, working at the intersection of network neuroscience and brain imaging, get started? A typical project involves myriad challenges that often encompass multiple established packages as well as custom code: interfacing with dataset APIs, implementing algorithms from a wide range of analytical traditions, and rendering brain surfaces to display results. Consider the task of community detection (identifying modules in brain connectomes): code to implement the quality function exists in several packages, but the wider framework to conduct the analysis—consensus clustering, mapping the space of degenerate partitions, etc.—does not. As a result, common practice is for groups to develop and informally share in-house scripts that rarely survive beyond one or two generations of trainees.

Here we present `netneurotools`, a Python utility for implementing exactly those tasks in network neuro-science for which there exists no straighforward solution. To our knowledge, `netneurotools` is a unique research product in the sense that it is the internal utility belt of a small lab. Initiated at the time of the lab’s inception at the Montréal Neurological Institute in 2017, the goal of the repository is simply to implement utilities that lab members frequently use. By design, functions are diverse and miscellaneous, including functions to construct network null models, estimate graph metrics, fit statistical models, fetch datasets, customize colormaps and plot brain images. The toolbox is “battle tested”: utilities have been extensively internally tested and used in > 50 publications. From the beginning, the repository has been open and has seen contributions from all over the world, including ≈ 30 forks and ≈ 75 stars on GitHub.

### OVERVIEW AND GUIDING PRINCIPLES

Given the broad use of the repository by our group and others, we have compiled a short report of what the current version of `netneurotools` entails and how it can be used (https://github.com/netneurolab/netneurotools). We emphasize that the utilities are intended to fill gaps between large, well-known software packages. Utilities are designed by trainees, for trainees: in this sense, the repository is an organic, living set of utilities that reflects the day-to-day needs of typical trainees working in the field. We close the report with several examples and an outlook for how the repository will continue to develop.

### Practical solutions for filling gaps in network neuroscience workflows

Researchers in network neuroscience are frequently challenged by seemingly “basic” routines that fall between the boundaries of major software packages. Questions such as how to import a brain atlas, parcellate brain maps, construct a robust consensus network, or flexibly visualize results on a brain surface are common yet surprisingly difficult to address with a single tool. Most importantly, researchers need solutions that utilize simple and straight-forward primitives without requiring them to learn new frameworks or data formats. Ideally, these solutions should generate reproducible results that can be readily integrated into existing analyses or workflows.

Despite the growing popularity of systems-level analysis, there are few actively maintained and time-tested toolboxes dedicated to addressing these practical challenges in network neuroscience. `netneurotools` fills this gap with a steadily growing collection of functions that aim at lowering barriers and standardizing atomic tasks in network neuroscience research. It builds on eight years of practical, data-intensive research inquiries by trainees who use these tools daily.

In this sense, the philosophy behind the toolbox is similar to the concept of “desire paths” in urban design. Here, paths emerge organically as individuals develop preferred routes through urban environments, such as by frequently walking through a park or courtyard. The repeated erosion caused by trampling is taken evidence of a desirable path and subsequently more formally implemented as a walkway by adding gravel or paving. `netneurotools` broadly follows the same idea of structure reflecting function: functionality and workflows are added only after we notice that they are frequently used by lab members.

### Toolbox architecture: accessibility and trainee-centric design

By design, `netneurotools` is trainee-centric and focuses on accessibility and usability. The functions are grouped into intuitive modules, covering the full lifecycle of a project (Fig. 1). A typical network neuroscience project commonly involves data ingestion, network reconstruction, metric estimation, statistical analysis, and visualization of results. A researcher starting a project can either bring their own data or simply proceed with the datasets module, where there is a large curated collection of standard templates, common atlases, and published datasets for fast and convenient prototyping. The interface module can handle potential incompatibilities from external data formats, tools, and modalities.

**Figure 1.**
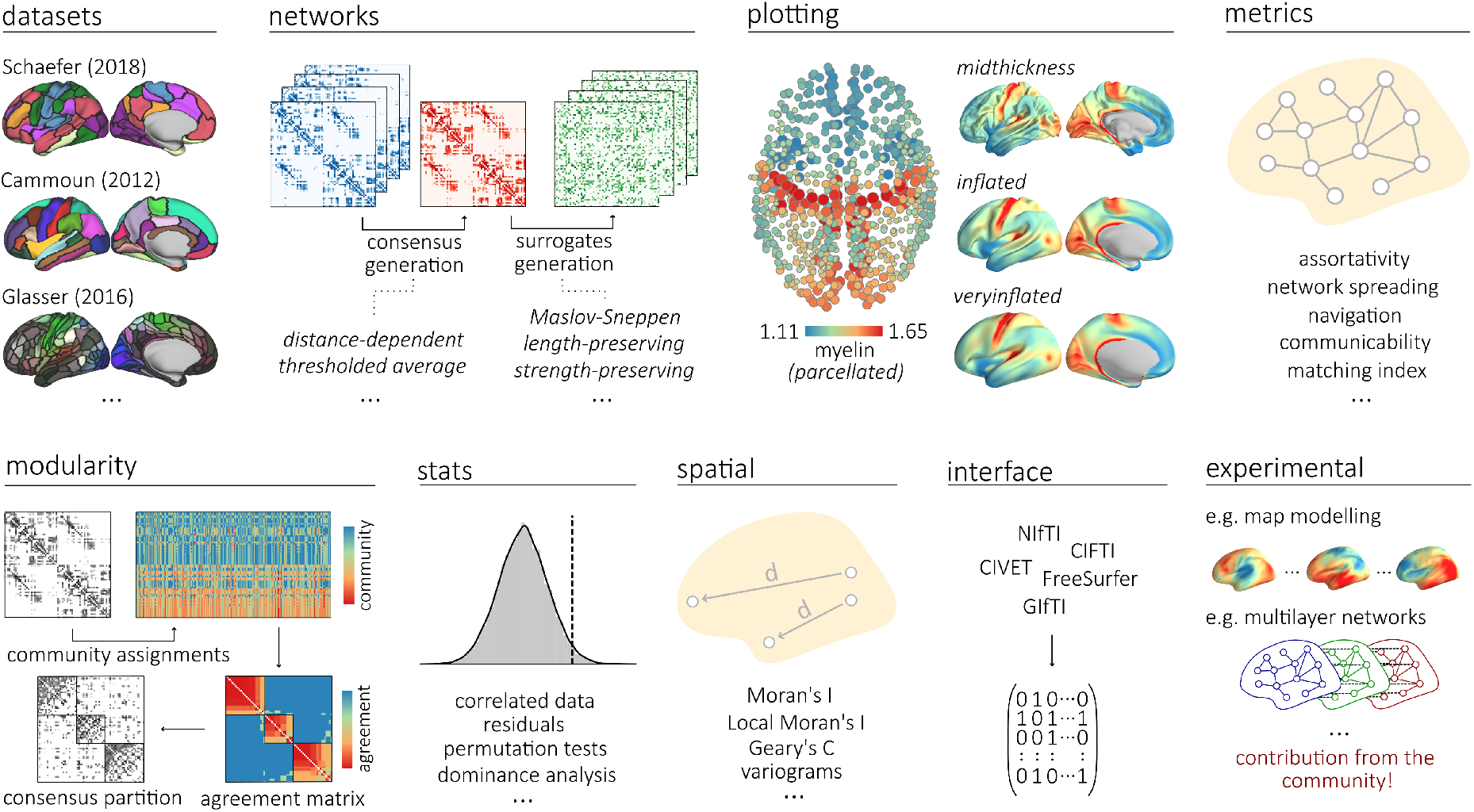
A Swiss army knife for network neuroscience. `netneurotools` implements a versatile and modular set of functions to support a wide range of analyses, from data preprocessing to network metrics and visualization. Functions are currently grouped in nine distinct modules: datasets, networks, plotting, metrics, modularity, stats, spatial, interface and experimental. See the *Modules and key functions* section for a brief overview of each module’s functionalities.

Following data ingestion, the networks module allows transforming raw connectivity data into several types of consensus and surrogate networks. The networks can then enter `metrics` and `modularity` modules, which provide optimized implementations of classic and cutting-edge network-theoretic metrics, including a streamlined community detection workflow. The stats module provides an extensive set of functions for correlations, regressions, and permutation tests. The final results can be visualized on the brain surface flexibly with the `plotting` module. By standardizing and streamlining fragmented computational routines into a curated toolbox, `netneurotools` lets trainees focus more on hypothesis discovery and reproducibility. It also stands as a high-quality, contextualized, and trustable reference of routine tasks in network neuroscience, compared to isolated code snippets online and generic responses from large language models.

From datasets to network metrics, from statistics to visualizations, the toolbox mirrors the steps trainees take daily. Adding to the broader usability, functions in netneurotools are modular and transparent, and are documented in great detail, allowing flexible mixing and matching. Trainees can parcellate their own data with curated atlases in `datasets`; pair `metrics` with external graph or machine learning libraries; use `stats` independently for permutation testing, etc.

The toolbox evolves organically and in tandem with the field—new modules and functions are added as researchers and the field specify new needs. For instance, the spatial module which implements geospatial statistics is the most recent addition towards studying `spatial` embedding and geometric features of the brain. An additional `experimental` module is available as a sandbox for testing new ideas from the community without disrupting the more stable utilities.

In Fig. 2, we showcase three examples of biologically-relevant questions that can be answered using `netneurotools` functionalities. These questions, inspired from previous work from our group [5, 22, 29], explore whether myelination levels are significantly more similar in structurally connected regions (Fig. 2a), whether structure-function coupling is consistent across functional modalities (Fig. 2b) and whether the brain’s community structure is consistent across modes of connectivity (Fig. 2c).

**Figure 2.**
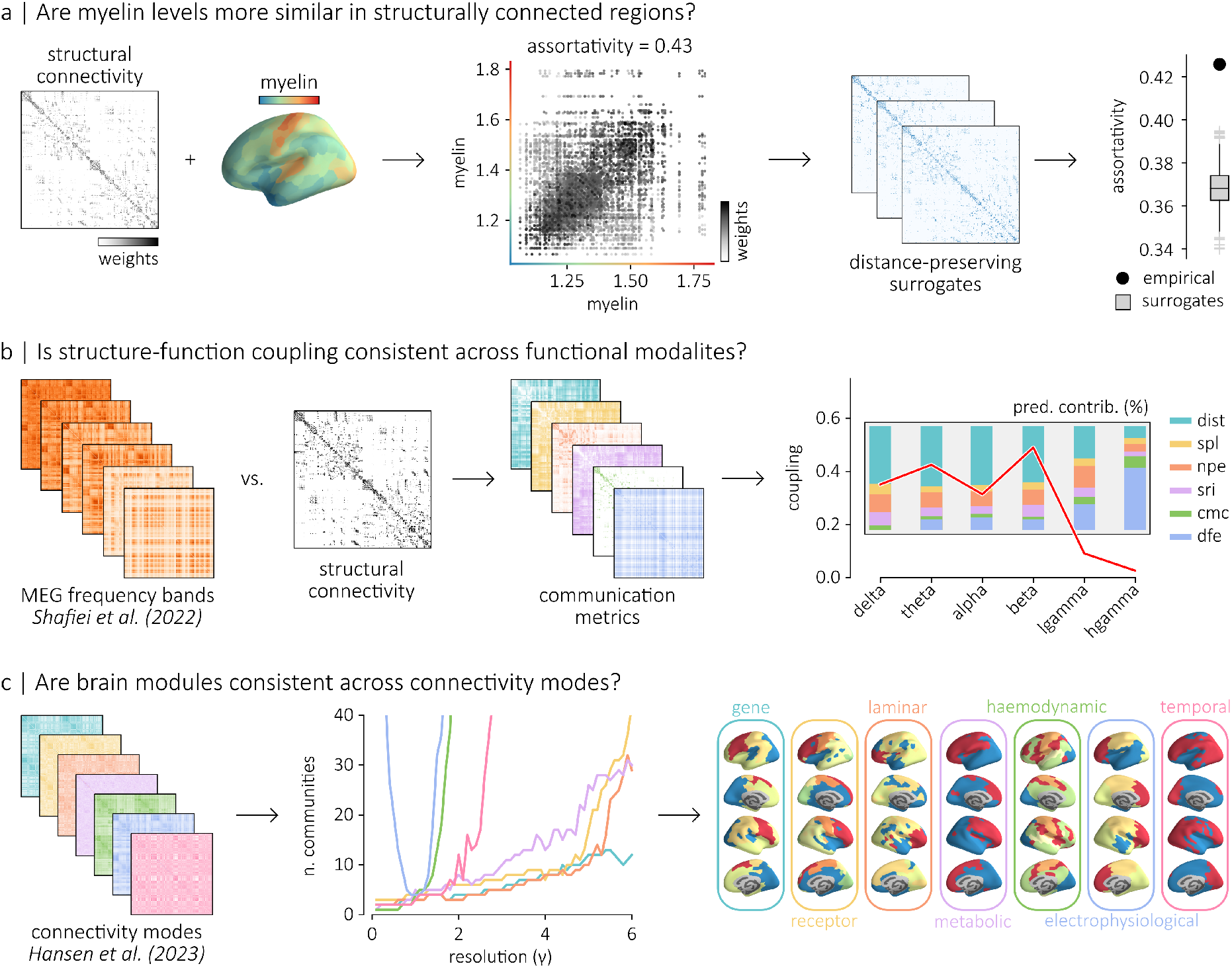
Answering neurobiologically meaningful questions with netneurotools. We provide three walkthrough examples for different stages of the research process and explicitly highlight functions that could be used at each stage. (a) In Bazinet et al. [5], we studied assortative mixing in annotated connectomes. The data used in this study can be directly downloaded from netneurotools using `fetch_bazinet_assortativity`. The assortativity of myelin with respect to the structural connectome can then be quantified using `assortativity_und`. To evaluate the significance of the assortativity coefficient, we can generate surrogate networks that preserve the edge length and degree distribution of the empirical networks with `match_length_degree_distribution`. In Liu et al. [29], we evaluated structure-function coupling across magnetoencephalography (MEG) frequency bands. We fetched MEG data from Shafiei et al. [41], using `fetch_shafiei_megfmrimapping`. Using netneurotools, we can compute communication metrics on the structural connectome (e.g. communicability with `communicability_wei`). We can then evaluate the coupling between MEG connectivity and structural connectivity with a multiple linear regression model using six communication metrics (dist: Euclidean distance, spl: shortest path length, npe: navigation efficiency, sri: search information, cmc: communicability, dfe: diffusion efficiency). To evaluate the contribution of each predictors, we can perform dominance analysis using `get_dominance_stats`. (c) In Hansen et al. [22], we compared the community structure of 7 distinct types of interregional similarity matrices (connectivity modes). The data can be downloaded directly from netneurotools using `fetch_hansen_manynetworks`. To identify the community structure of each connectivity mode for a given resolution *γ*, we can use `consensus_modularity`, which performs multiple iterations of community detection using the Louvain algorithm [7] and then uses clustering to derive a consensus community assignment, as detailed in Bassett et al. [4]. Finally, the community assignments can be visualized on the surface of the cortex using `plot_fsaverage`.

### Sustainable development and community building

Maintaining scientific software requires concerted effort, especially while supporting an active user base alongside academic commitments [48]. In the process of keeping `netneurotools` relevant and publication-ready over time, we have learned several important principles. First, we maintain minimal dependencies and broad applicability. `netneurotools` is embedded in the Python scientific computing ecosystem, directly built on fundamental packages including those for numerical computing (NumPy [23], SciPy [46], Scikit-learn [34], NetworkX [21]), visualization (Matplotlib [24]), and quantitative neuroimaging (NiBabel [8], Nilearn [33]). This foundation ensures stability and compatibility with all major platforms. Users can optionally install additional dependencies depending on specific use cases for speed-optimized functions (Numba [25], Joblib [43]) and cortical surface rendering (VTK [39], PyVista [42]).

Second, we prioritize automated, reproducible builds. For every update and versioned release, `netneurotools` is automatically built with GitHub Actions as Docker containers. Updates are automatically tested to ensure that core functionality remains reliable. A continuously updated documentation is similarly compiled and hosted on GitHub Pages (https://netneurolab.github.io/netneurotools/), providing detailed function references, examples, and tutorials.

Third, we encourage sustainable community involvement both internally and externally. Within our lab, trainees are encouraged to reuse and improve the existing functions, as well as incorporating new functions from their projects in `netneurotools`. Externally, we actively collaborate with researchers worldwide to incorporate new functions and address community needs through a support ticket system. The research-driven development ensures that the toolbox stays updated with the frontier questions and methods in network neuroscience and brain imaging. We also actively foster an open science environment to enable continual transfer of knowledge transcending the typical graduate student lifecycle. Functions in `netneurotools` originally developed by early lab members are now being maintained and extended by newer generations of trainees.

## MODULES AND KEY FUNCTIONS

### The netneurotools package

The `netneurotools` package is organized into 9 modules that provide comprehensive functionality for data acquisition, network construction, graph theory calculations, visualization, statistical analysis, and interfacing with external neuroimaging tools. `netneurotools` is distributed under the BSD-3 license, making it free for academic and commercial use. The package is available on GitHub (https://github.com/netneurolab/netneurotools) with comprehensive documentation (https://netneurolab.github.io/netneurotools/). It can be installed via PyPI (pip install `netneurotools`), from source, or used from a Docker container.

### Toolbox dependencies

`netneurotools` depends on a minimal set of scientific Python packages: NumPy [23], SciPy [46], Scikit-learn [34], and NetworkX [21] for numerical procedures; NiBabel [8] and Nilearn [33] for routine neuroimaging operations. The package leverages Joblib [43] for parallelization and Numba [25] for just-in-time compilation of computationally intensive functions. For visualization, `netneurotools` utilizes Matplotlib [24] for 2D figures and PyVista [42], a streamlined interface to the Visualization Toolkit (VTK) [39], for publication-quality 3D renderings of brain surfaces.

### Ingesting data: the datasets and networks modules

The `netneurotools.datasets` module provides an integrated infrastructure for programmatically downloading, loading, and managing various neuroimaging datasets, atlases, templates, and published results. Key functions include fetching cortical surface templates (e.g., `fetch_fsaverage` [17], `fetch_conte69` [44]), brain parcellations (e.g., `fetch_cammoun2012` [10], `fetch_schaefer2018` [38]), and published datasets (e.g., `fetch_vazquez_rodriguez2019` [45]). The module implements a caching system to efficiently store and retrieve downloaded data. The `netneurotools.networks` module contains methods for generating and preprocessing brain networks. It includes functions for creating consensus functional (`func_consensus`) and structural (`struct_consensus` [6]) connectivity matrices from a group of subjects, generating randomized networks while preserving desired topological or geometrical properties (`randmio_und` [30], `strength_preserving_rand_sa` [31]), and thresholding networks (`threshold_network, binarize_network`).

### Visualizing the brain: the plotting module

The `netneurotools.plotting` module offers routines for creating publication-ready visualizations of volumetric, surface, and network-structured brain data. Surface-based plotting functions (`plot_fsaverage` and `pv_plot_surface`) allow convenient and flexible mapping of cortical surfaces, with customizable views, lights, and colors. Coordinate-based plotting function (`plot_point_brain`) renders brain regions as 3D point clouds. This module also provides specialized functions for visualizing network communities (`plot_mod_heatmap`).

### Calculating graph metrics: the metrics and modularity modules

The netneurotools.metrics module features an optimized collection of functions for working with brain networks, implementing both classical and novel graph theoretical measures [37]. These include functions for calculating basic nodal properties (`degrees_und, degrees_dir`), shortest path and routing metrics (`distance_wei_floyd` [18, 36, 47], `navigation_wu` [40]), communication dynamics (`communicability_bin` [15], `communicability_wei` [13], `mean_first_passage_time`) and statistical network metrics (`network_pearsonr` [12], `network_variance` [14]). Many implementations offer significant performance improvements over existing packages through vectorization and Numba acceleration.

The `netneurotools.modularity` module includes functions for exploring and analyzing community structure in brain networks. It provides tools for identifying consensus community assignments across multiple partitioning solutions (`find_consensus` [4]), assessing the quality of community assignments (`get_modularity, get_modularity_z`), and calculating partition similarity (`zrand` [35]).

### Estimating spatially naive and spatially embedded statistics: the stats and spatial modules

The `netneurotools.stats` module provides functions for statistical analysis and inference in neuroimaging. It includes efficient implementations of correlation metrics (`efficient_pearsonr, weighted_pearsonr`), permutation testing routines (`permtest_1samp, permtest_pearsonr`), and regression analyses (`residualize, get_dominance_stats` [2, 9]).

The `netneurotools.spatial` module features functions for studying the spatial organization of neural patterns, offering tools to calculate spatial autocorrelation metrics (Moran’s I [11, 28, 32], Geary’s C [1, 19], Lee’s L [27]) and their local variants. These functions enable researchers to study the spatial dependence of structural and functional brain features.

### Navigating complex tools: the interface module

The `netneurotools.interface` module includes functions for interacting with external neuroimaging tools and file formats. It provides parsers and converters for FreeSurfer [16] (`extract_annot_labels`), CIFTI [20] (`extract_cifti_surface`), and GIfTI formats, facilitating the extraction and manipulation of data from these complex file formats, enabling seamless workflow integration with established neuroimaging software packages.

### Incubating new ideas: the experimental module

The `netneurotools.experimental` module allows users to experiment with cutting-edge methods before they are fully integrated into the main package. It is a sandbox and incubator to try out functions that are novel but still unstable. It thus facilitates and encourages new contributions from the community which could include, for example, new methods for multilayer network analysis or for surrogate brain map generation.

### Automated testing and release

`netneurotools` employs continuous integration through GitHub Actions to ensure code quality and reliability. The automated process runs comprehensive unit tests across multiple Python versions and operating systems upon code changes, ensuring that new contributions maintain compatibility and functionality. The package is regularly released and published to PyPI and Docker Hub, making it readily available to the scientific community.

## CONCLUDING REMARKS

The challenge of accessible and reproducible research practices is substantial and ongoing. `netneurotools` brings not only tools of the trade but also embodies our efforts of implementing FAIR principles (Findable, Accessible, Interoperable, and Reusable) [49] in research coding [3, 26]. By curating standardized procedures, facilitating data sharing, and ensuring code transparency, we hope `netneurotools` lower the barrier and raise the rigor in the field of network neuroscience.

As the field continues to grow, so too will `netneurotools`. We envision it as an evolving set of utilities that reflect the day-to-day needs of both new and experienced trainees. Through research-driven, sustainable development, we aim to ensure that `netneurotools` remains a useful resource for the community: bridging gaps between specialized tools and empowering researchers to focus on scientific discovery rather than technical implementation.

## ACKNOWLEDGMENTS

We thank Yigu Zhou, Moohebat Pourmajidian, and Tahmineh Taheri for comments and suggestions on the manuscript. We also thank past and future contributors to the toolbox. BM acknowledges support from the Natural Sciences and Engineering Research Council of Canada (NSERC), Canadian Institutes of Health Research (CIHR), Brain Canada Foundation Future Leaders Fund, the Canada Research Chairs Program, the Michael J. Fox Foundation, and the Healthy Brains for Healthy Lives initiative. ZQL acknowledges support from the Fonds de Recherche du Québec – Nature et Technologies (FRQNT). VB acknowledges support from the Natural Sciences and Engineering Research Council of Canada (NSERC), the Fonds de Recherche du Quebec - Nature et Technologie (FRQNT), the Healthy Brains for Healthy Lives initiative and Brain Canada.

## References

[1] Anselin, L. (2019). A Local Indicator of Multivariate Spatial Association: Extending Geary’s c. Geographical Analysis, 51(2):133–150.

[2] Azen, R. and Budescu, D. V. (2003). The dominance analysis approach for comparing predictors in multiple regression. Psychological Methods, 8(2):129–148.

[3] Barker, M., Chue Hong, N. P., Katz, D. S., Lamprecht, A.-L., Martinez-Ortiz, C., Psomopoulos, F., Harrow, J., Castro, L. J., Gruenpeter, M., Martinez, P. A., and Honeyman, T. (2022). Introducing the FAIR Principles for research software. Scientific Data, 9(1):622.

[4] Bassett, D. S., Porter, M. A., Wymbs, N. F., Grafton, S. T., Carlson, J. M., and Mucha, P. J. (2013). Robust detection of dynamic community structure in networks. Chaos: An Interdisciplinary Journal of Nonlinear Science, 23(1):013142.

[5] Bazinet, V., Hansen, J. Y., Vos de Wael, R., Bernhardt, B. C., van den Heuvel, M. P., and Misic, B. (2023). Assortative mixing in micro-architecturally annotated brain connectomes. Nature Communications, 14(1):2850.

[6] Betzel, R. F., Griffa, A., Hagmann, P., and Mišić, B. (2018). Distance-dependent consensus thresholds for generating group-representative structural brain networks. Network Neuroscience, 3(2):475–496.

[7] Blondel, V. D., Guillaume, J.-L., Lambiotte, R., and Lefebvre, E. (2008). Fast unfolding of communities in large networks. Journal of Statistical Mechanics: Theory and Experiment, 2008(10):P10008.

[8] Brett, M., Markiewicz, C. J., Hanke, M., Côté, M.-A., Cipollini, B., Papadopoulos Orfanos, D., McCarthy, P., Jarecka, D., Cheng, C. P., Larson, E., Halchenko, Y. O., Cottaar, M., Ghosh, S., Wassermann, D., Gerhard, S., Lee, G. R., Baratz, Z., Moloney, B., Wang, H.-T., Kastman, E., Kaczmarzyk, J., Guidotti, R., Daniel, J., Duek, O., Rokem, A., Scheltienne, M., Madison, C., Sólon, A., Morency, F. C., Goncalves, M., Markello, R., Riddell, C., Burns, C., Millman, J., Gramfort, A., Leppäkangas, J., van den Bosch, J. J., Vincent, R. D., Braun, H., Subramaniam, K., Van, A., Legarreta, J. H., Gorgolewski, K. J., Raamana, P. R., Klug, J., Vos de Wael, R., Nichols, B. N., Baker, E. M., Koudoro, S., Hayashi, S., Pinsard, B., Haselgrove, C., Hymers, M., Esteban, O., Pérez-García, F., Becq, G., Dockès, J., Oosterhof, N. N., Amirbekian, B., Christian, H., Nimmo-Smith, I., Nguyen, L., Suter, P., Reddigari, S., St-Jean, S., Panfilov, E., Garyfallidis, E., Varoquaux, G., Newton, J., Hahn, K. S., Waller, L., Hinds, O. P., Sandro Fauber, B., Dewey, B., Perez, F., Roberts, J., Poline, J.-B., Stutters, J., Jordan, K., Cieslak, M., Moreno, M. E., Hrnčiar, T., Haenel, V., Schwartz, Y., Darwin, B. C., Thirion, B., Gauthier, C., Solovey, I., Gonzalez, I., Palasubramaniam, J., Lecher, J., Leinweber, K., Raktivan, K., Calábková, M., Fischer, P., Gervais, P., Gadde, S., Ballinger, T., Roos, T., Reddam, V. R., and freec84 (2024). Nipy/nibabel. Zenodo.

[9] Budescu, D. V. (1993). Dominance analysis: A new approach to the problem of relative importance of predictors in multiple regression. Psychological Bulletin, 114(3):542–551.

[10] Cammoun, L., Gigandet, X., Meskaldji, D., Thiran, J. P., Sporns, O., Do, K. Q., Maeder, P., Meuli, R., and Hagmann, P. (2012). Mapping the human connectome at multiple scales with diffusion spectrum MRI. Journal of Neuroscience Methods, 203(2):386–397.

[11] Cliff, A. D. and Ord, J. K. (1981). Spatial Processes: Models & Applications. Pion.

[12] Coscia, M. (2021). Pearson correlations on complex networks. Journal of Complex Networks, 9(6):cnab036.

[13] Crofts, J. J. and Higham, D. J. (2009). A weighted communicability measure applied to complex brain networks. Journal of The Royal Society Interface, 6(33):411–414.

[14] Devriendt, K., Martin-Gutierrez, S., and Lambiotte, R. (2022). Variance and Covariance of Distributions on Graphs. SIAM Review, 64(2):343–359.

[15] Estrada, E. and Hatano, N. (2008). Communicability in complex networks. Physical Review E, 77(3):036111.

[16] Fischl, B. (2012). FreeSurfer. NeuroImage, 62(2):774– 781.

[17] Fischl, B., Sereno, M. I., Tootell, R. B., and Dale, A. M. (1999). High-resolution intersubject averaging and a coordinate system for the cortical surface. Human Brain Mapping, 8(4):272–284.

[18] Floyd, R. W. (1962). Algorithm 97: Shortest path. Communications of the ACM, 5(6):345–345.

[19] Geary, R. C. (1954). The Contiguity Ratio and Statistical Mapping. The Incorporated Statistician, 5(3):115.

[20] Glasser, M. F., Sotiropoulos, S. N., Wilson, J. A., Coalson, T. S., Fischl, B., Andersson, J. L., Xu, J., Jbabdi, S., Webster, M., Polimeni, J. R., Van Essen, D. C., and Jenkinson, M. (2013). The minimal preprocessing pipelines for the Human Connectome Project. NeuroImage, 80:105–124.

[21] Hagberg, A. A., Schult, D. A., and Swart, P. J. (2008). Exploring Network Structure, Dynamics, and Function using NetworkX. In Proceedings of the Python in Science Conference, pages 11–15. SciPy.

[22] Hansen, J. Y., Shafiei, G., Voigt, K., Liang, E. X., Cox, S. M. L., Leyton, M., Jamadar, S. D., and Misic, B. (2023). Integrating multimodal and multiscale connectivity blueprints of the human cerebral cortex in health and disease. PLOS Biology, 21(9):e3002314.

[23] Harris, C. R., Millman, K. J., van der Walt, S. J., Gommers, R., Virtanen, P., Cournapeau, D., Wieser, E., Taylor, J., Berg, S., Smith, N. J., Kern, R., Picus, M., Hoyer, S., van Kerkwijk, M. H., Brett, M., Haldane, A., del Río, J. F., Wiebe, M., Peterson, P., Gérard-Marchant, P., Sheppard, K., Reddy, T., Weckesser, W., Abbasi, H., Gohlke, C., and Oliphant, T. E. (2020). Array programming with NumPy. Nature, 585(7825):357–362.

[24] Hunter, J. D. (2007). Matplotlib: A 2D Graphics Environment. Computing in Science & Engineering, 9(3):90–95.

[25] Lam, S. K., Pitrou, A., and Seibert, S. (2015). Numba: A LLVM-based Python JIT compiler. In Proceedings of the Second Workshop on the LLVM Compiler Infrastructure in HPC, LLVM ‘15, pages 1–6, New York, NY, USA. Association for Computing Machinery.

[26] Lamprecht, A.-L., Garcia, L., Kuzak, M., Martinez, C., Arcila, R., Martin Del Pico, E., Dominguez Del Angel, V., van de Sandt, S., Ison, J., Martinez, P. A., McQuilton, P., Valencia, A., Harrow, J., Psomopoulos, F., Gelpi, J. L., Chue Hong, N., Goble, C., and Capella-Gutierrez, S. (2020). Towards FAIR principles for research software. Data Science, 3(1):37–59.

[27] Lee, S.-I. (2001). Developing a bivariate spatial association measure: An integration of Pearson’s r and Moran’s I. Journal of Geographical Systems, 3(4):369–385.

[28] Li, H., Calder, C. A., and Cressie, N. (2007). Beyond Moran’s I: Testing for Spatial Dependence Based on the Spatial Autoregressive Model. Geographical Analysis, 39(4):357–375.

[29] Liu, Z.-Q., Shafiei, G., Baillet, S., and Misic, B. (2023). Spatially heterogeneous structure-function coupling in haemodynamic and electromagnetic brain networks. NeuroImage, 278:120276.

[30] Maslov, S. and Sneppen, K. (2002). Specificity and Stability in Topology of Protein Networks. Science, 296(5569):910–913.

[31] Milisav, F., Bazinet, V., Betzel, R. F., and Misic, B. (2025). A simulated annealing algorithm for randomizing weighted networks. Nature Computational Science, 5(1):48–64.

[32] Moran, P. A. P. (1950). Notes on Continuous Stochastic Phenomena. Biometrika, 37(1/2):17.

[33] Nilearn contributors, Chamma, A., Frau-Pascual, A., Rothberg, A., Abadie, A., Abraham, A., Gramfort, A., Savio, A., Cionca, A., Sayal, A., Thual, A., Kodibagkar, A., Kanaan, A., Pinho, A. L., Joshi, A., Idrobo, A. H., Kieslinger, A.-S., Kumari, A., Rokem, A., Mensch, A., Vijayan, A., Duran, A., Cipollini, B., Thirion, B., Nguyen, B., Cakan, C., Gorgolewski, C., Markiewicz, C., Horea, C., Gerloff, C., Roßmanith, C., Reininger, C., Lane, C., Sy, C., Delettre, C., Gale, D., Gomez, D., Bzdok, D., Ellis, D. G., Wassermann, D., Pisner, D., Orfanos, D. P., DuPre, E., Dohmatob, E., Larson, E., Edmond, E., Pedregosa, F., Pollet, F., Liem, F., Paugam, F., Varoquaux, G., de Hollander, G., Kiar, G., Gilmore, G., Lemaitre, G., Gözükan, H., Wang, H.-T., Aggarwal, H., Abenes, I., Traore, I., Vogel, J., Margeta, J., Grobler, J., Gors, J., Kai, J., Rasero, J., Kossaifi, J., King, J.-R., Dalenberg, J. R., Lefort-Besnard, J., Dockes, J., Chevalier, J.-A., Wiesner, J., Johnson, J. T., Gorrono, J. H. L., Sassenhagen, J., Huguet, J., Teves, J., Huntenburg, J., Peraza, J. A., Daddy, K. R., Sitek, K., Helwegen, K., Wagstyl, K., Shmelkov, K., Chawla, K., Chen, K., Newberg, L., Sasse, L., Estève, L., Tetrel, L., Paz, L., Pietrantoni, M., Perez-Guevara, M., Wegrzyn, M., Goncalves, M., Dugré, M., Ekman, M., Joulot, M., Sitter, M. C., Rahim, M., Zwally, M., Burkhardt, M., Eickenberg, M., Hanke, M., Notter, M., Waskom, M., Wang, M., Torabi, M., Boos, M., Chapra, M., Song, M. S., Clarke, N., Shah, N., Gensollen, N., Krish, N., Warrington, O., Esteban, O., Sadil, P., Bogdan, P., Reiners, P., Sanz-Leon, P., Herholz, P., Gervais, P., Bellec, P., Glaser, P., Barbarant, P.-L., Quirion, P.-O., Raamana, P. R., Jain, P., Brito, R., Meudec, R., Luke, R., Williamson, R., Guidotti, R., Jepegnanam, R. T., Phlypo, R., Hammonds, R., Gau, R., Patalasingh, S., Hahn, S., Bougacha, S., Johnson, S. B., Jawhar, S., Steinkamp, S., Kim, S., Singh, S., Meisler, S., Pokharel, S., Lan, S., Takerkart, S., Gezici, T., Samanta, T., Salo, T. K. T., Premrudeepreechacharn, T., Bazeille, T., Vanasse, T., Diogo, V., Shevchenko, V., Michel, V., Fritsch, V., Halchenko, Y., Mzayek, Y., Huang, Y., Baratz, Z., and Nájera, Ó. (2025). Nilearn. Zenodo.

[34] Pedregosa, F., Varoquaux, G., Gramfort, A., Michel, V., Thirion, B., Grisel, O., Blondel, M., Prettenhofer, P., Weiss, R., Dubourg, V., Vanderplas, J., Passos, A., Cournapeau, D., Brucher, M., Perrot, M., and Duchesnay, É. (2011). Scikit-learn: Machine Learning in Python. Journal of Machine Learning Research, 12(85):2825–2830.

[35] Red, V., Kelsic, E. D., Mucha, P. J., and Porter, M. A. (2011). Comparing Community Structure to Characteristics in Online Collegiate Social Networks. SIAM Review, 53(3):526–543.

[36] Roy, B. (1959). Transitivité et connexité. CR Acad. Sci. Paris, 249(216–218):182.

[37] Rubinov, M. and Sporns, O. (2010). Complex network measures of brain connectivity: Uses and interpretations. NeuroImage, 52(3):1059–1069.

[38] Schaefer, A., Kong, R., Gordon, E. M., Laumann, T. O., Zuo, X.-N., Holmes, A. J., Eickhoff, S. B., and Yeo, B. T. T. (2018). Local-Global Parcellation of the Human Cerebral Cortex from Intrinsic Functional Connectivity MRI. Cerebral Cortex, 28(9):3095–3114.

[39] Schroeder, W., Martin, K. M., and Lorensen, W. E. (1998). The Visualization Toolkit an Object-Oriented Approach to 3D Graphics. Prentice-Hall, Inc.

[40] Seguin, C., van den Heuvel, M. P., and Zalesky, A. (2018). Navigation of brain networks. Proceedings of the National Academy of Sciences, 115(24):6297–6302.

[41] Shafiei, G., Baillet, S., and Misic, B. (2022). Human electromagnetic and haemodynamic networks systematically converge in unimodal cortex and diverge in transmodal cortex. PLOS Biology, 20(8):e3001735.

[42] Sullivan, C. and Kaszynski, A. (2019). PyVista: 3D plotting and mesh analysis through a streamlined interface for the Visualization Toolkit (VTK). Journal of Open Source Software, 4(37):1450.

[43] The joblib developers (2025). Joblib. Zenodo.

[44] Van Essen, D. C., Glasser, M. F., Dierker, D. L., Harwell, J., and Coalson, T. (2012). Parcellations and Hemispheric Asymmetries of Human Cerebral Cortex Analyzed on Surface-Based Atlases. Cerebral Cortex, 22(10):2241– 2262.

[45] Vázquez-Rodríguez, B., Suárez, L. E., Markello, R. D., Shafiei, G., Paquola, C., Hagmann, P., van den Heuvel, M. P., Bernhardt, B. C., Spreng, R. N., and Misic, B. (2019). Gradients of structure–function tethering across neocortex. Proceedings of the National Academy of Sciences, 116(42):21219–21227.

[46] Virtanen, P., Gommers, R., Oliphant, T. E., Haberland, M., Reddy, T., Cournapeau, D., Burovski, E., Peterson, P., Weckesser, W., Bright, J., van der Walt, S. J., Brett, M., Wilson, J., Millman, K. J., Mayorov, N., Nelson, A. R. J., Jones, E., Kern, R., Larson, E., Carey, C. J., Polat, İ., Feng, Y., Moore, E. W., VanderPlas, J., Laxalde, D., Perktold, J., Cimrman, R., Henriksen, I., Quintero, E. A., Harris, C. R., Archibald, A. M., Ribeiro, A. H., Pedregosa, F., and van Mulbregt, P. (2020). SciPy 1.0: Fundamental algorithms for scientific computing in Python. Nature Methods, 17(3):261–272.

[47] Warshall, S. (1962). A Theorem on Boolean Matrices. Journal of the ACM, 9(1):11–12.

[48] Westner, B. U., McCloy, D. R., Larson, E., Gramfort, A., Katz, D. S., Smith, A. M., Delorme, A., Litvak, V., Makeig, S., Oostenveld, R., Schoffelen, J.-M., and Tierney, T. M. (2025). Cycling on the Freeway: The perilous state of open-source neuroscience software. Imaging Neuroscience, 3:imag_a_00554.

[49] Wilkinson, M. D., Dumontier, M., Aalbersberg, I. J., Appleton, G., Axton, M., Baak, A., Blomberg, N., Boiten, J.-W., Santos, L. B. d. S., Bourne, P. E., Bouwman, J., Brookes, A. J., Clark, T., Crosas, M., Dillo, I., Dumon, O., Edmunds, S., Evelo, C. T., Finkers, R., Gonzalez-Beltran, A., Gray, A. J. G., Groth, P., Goble, C., Grethe, J. S., Heringa, J., ‘t Hoen, P. A. C., Hooft, R., Kuhn, T., Kok, R., Kok, J., Lusher, S. J., Martone, M. E., Mons, A., Packer, A. L., Persson, B., Rocca-Serra, P., Roos, M., van Schaik, R., Sansone, S.-A., Schultes, E., Sengstag, T., Slater, T., Strawn, G., Swertz, M. A., Thompson, M., van der Lei, J., van Mulligen, E., Velterop, J., Waagmeester, A., Wittenburg, P., Wolstencroft, K., Zhao, J., and Mons, B. (2016). The FAIR Guiding Principles for scientific data management and stewardship. Scientific Data.

